# An 18S V4 rDNA metabarcoding dataset of protist diversity in the Atlantic inflow to the Arctic Ocean, through the year and down to 1000 m depth

**DOI:** 10.1101/2021.03.29.437475

**Authors:** Elianne Egge, Stephanie Elferink, Daniel Vaulot, Uwe John, Gunnar Bratbak, Aud Larsen, Bente Edvardsen

## Abstract

Arctic marine protist communities have been understudied due to challenging sampling conditions, in particular during winter and in deep waters. The aim of this study was to improve our knowledge on Arctic protist diversity through the year, both in the epipelagic (< 200 m depth) and mesopelagic zones (200-1000 m depth). Sampling campaigns were performed in 2014, during five different months, to capture the various phases of the Arctic primary production: January (winter), March (pre-bloom), May (spring bloom), August (post-bloom) and November (early winter). The cruises were undertaken west and north of the Svalbard archipelago, where warmer Atlantic waters from the West Spitsbergen Current meets cold Arctic waters from the Arctic Ocean. From each cruise, station, and depth, 50 L of sea water were collected and the plankton was size-fractionated by serial filtration into four size fractions between 0.45-200 *µ*m, representing the picoplankton, nanoplankton and microplankton. In addition vertical net hauls were taken from 50 m depth to the surface at selected stations. From the plankton samples DNA was extracted, the V4 region of the 18S rRNA-gene was amplified by PCR with universal eukaryote primers and the amplicons were sequenced by Illumina high-throughput sequencing. Sequences were clustered into Amplicon Sequence Variants (ASVs), representing protist genotypes, with the dada2 pipeline. Taxonomic classification was made against the curated Protist Ribosomal Reference database (PR2). Altogether 6,536 protist ASVs were obtained (including 54 fungal ASVs). Both ASV richness and taxonomic composition were strongly dependent on size-fraction, season, and depth. ASV richness was generally higher in the smaller fractions, and higher in winter and the mesopelagic samples than in samples from the well-lit epipelagic zone during summer. During spring and summer, the phytoplankton groups diatoms, chlorophytes and haptophytes dominated in the epipelagic zone. Parasitic and heterotrophic groups such as Syndiniales and certain dinoflagel-lates dominated in the mesopelagic zone all year, as well as in the epipelagic zone during the winter. The dataset is available at https://doi.org/10.17882/79823, (Egge et al., 2014).

## 1 Introduction

The West Spitsbergen current is considered the main gateway from the Atlantic into the Arctic Ocean, as it flows along the west side of the Svalbard Archipelago, transporting relatively warm and salty water (T > 2*°*C, S > 34.92; c.f. Randelhoff et al. (2018)) into the Barents Sea and Arctic Ocean (Figure 1). In response to global warming, this current has become both warmer and stronger in recent years, increasingly replacing water advected from the central Arctic Ocean with warm and salty water of Atlantic origin, a process referred to as “Atlantification” (Årthun et al., 2012). This increase in oceanic heat in the Arctic area correlates with the rapid decline in ice extent observed over the past decades (Årthun et al., 2012). Increased inflow of Atlantic water affects the primary production and protist communities in several ways. Water mixing happens more easily in the Atlantic water, because in contrast to the permanently salinity-stratified central Arctic Ocean, the water column is temperature-stratified and less stable, thus upper-ocean nutrients are more efficiently replenished early in winter. Furthermore, the warm Atlantic water melts the ice, and a layer of fresh, cold water is formed near the surface. The timing of this stratification is crucial for the onset of the spring bloom, and thinner ice means less light-limitations for the algae living inside and under the ice. However, the loss of sea ice also results in the loss of habitat for many protists, especially those adapted to a life in or on the ice. These various effects of climate change may thus alter both the location and timing of blooms, as well as their biomass and species composition (e.g. Eamer et al., 2013; Li et al., 2009).

**Figure 1.**
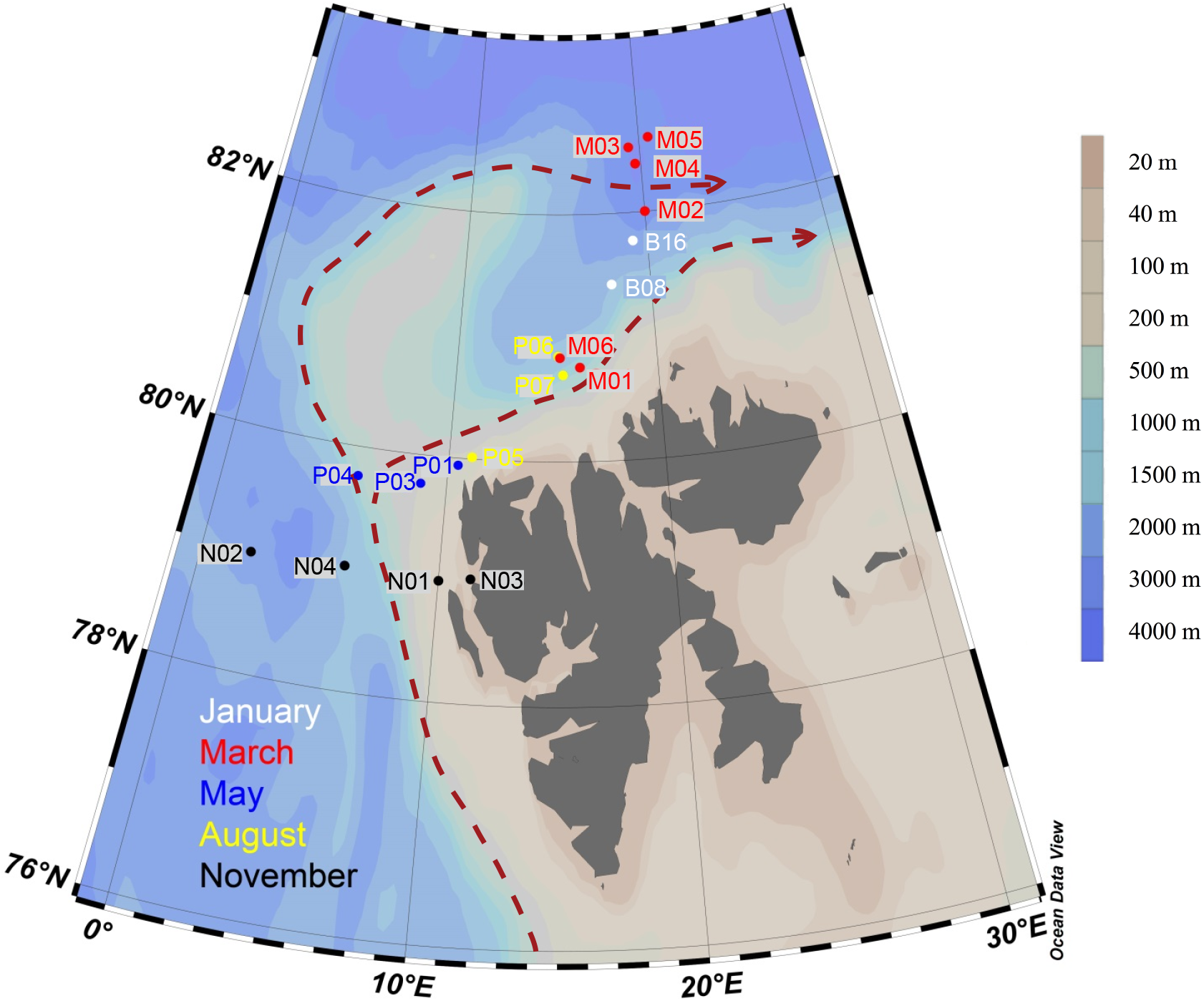
Map of sampling locations of the MicroPolar sampling campaign. Color correspond to cruise month. The red dashed line indicates the major flow patterns of warm Atlantic Water into the Arctic Ocean. Color scale bar indicates bottom depth. Red arrows indicate the main flow of the West Spitsbergen Current, according to Cokelet et al. (2008) and Randelhoff et al. (2018).

To understand what consequences environmental changes in this Arctic region will have for the biodiversity of the whole pelagic community and for the production through the food web up to higher trophic levels, we need to know what are the community components and where and how the organisms occur. This will also enable us to detect future changes. However, still relatively little is known about the diversity and distribution of protists in the Arctic Ocean (e.g. Lovejoy, 2014). Microbial eukaryote communities during the winter season and at mesopelagic depths (i.e. below the photic zone, 200-1000 m depth) are particularly understudied due to logistic challenges. Metabarcoding using high-throughput sequencing (HTS) has become a commonly used method to study the community composition of marine protists, and has revealed a huge unknown diversity (e.g. de Vargas et al., 2015). In recent years, several metabarcoding studies of protist communities in the Arctic Ocean have been undertaken, but most represent only snapshots of the community as based on a single cruise or season (e.g. Bachy et al., 2011; Kilias et al., 2014; Monier et al., 2015; Vader et al., 2015). Studies that have sampled the full yearly cycle have typically only sampled the upper water column (0-50m depth) (e.g. Marquardt et al., 2016).

Here we present a metabarcoding dataset from the Northern Svalbard region of the Arctic Ocean sampled during five cruises representing the full seasonal cycle, and at 3-4 depths from the surface down to 1000 m. Metabarcoding targeted the V4 region of the 18S rRNA gene. The data are provided both as raw reads and as Amplicon Sequence Variants obtained after processing with the dada2 pipeline, with corresponding ASV abundance tables. The data presented here were obtained within the framework of the project ‘MicroPolar’ (https://www.researchinsvalbard.no/project/7280). The virus and prokaryote communities from the same project have been described in Sandaa et al. (2018), and Wilson et al. (2017) and Paulsen et al. (2016), respectively. Environmental data from the MicroPolar sampling campaign have previously been published in Paulsen et al. (2017) and Randelhoff et al. (2018). A subset of the environmental data corresponding to the stations and depths of the protist metabarcoding samples is included in the data repository of the present study.

## 2 Study area and general environmental conditions

### 2.1 Study area

Sampling campaigns were performed in 2014 as described in detail in Paulsen et al. (2017), during five different months, to capture the various stages of the Arctic primary production: January (06.01–15.01, winter), March (05.03-10.03, pre-bloom), May (15.05-02.06, spring bloom), August (07.08-18.08, post-bloom) and November (03.11-10.11, early winter). The cruises were undertaken west and north of the Svalbard archipelago, where warmer Atlantic water in the West Spitsbergen Current meets colder water from the Arctic Ocean (Figure 1). Bottom depth varied from 327 m (November station N03) to c. 3000 m (March station M05). The area and locations for each sampling campaign were as similar as possible, but constrained by the sea ice cover, from 79 - 82.6 *°*N. During each cruise, transects of 3-6 stations were sampled at three or four depths: in the epipelagic zone at 1 m and at the deep chlorophyll maximum (usually between 15-25 m), and in the mesopelagic zone at one or two depths, as a rule 500 m and 1000 m, or as deep as the bathymetry of the station permitted. The ice extent was smallest in January, and peaked in May (see Figure 1 in Wilson et al., 2017). The two stations sampled in January were in the open ocean, whereas in March, May and August, all the stations were situated in varying degrees of drift ice, except March station M06 and August station P05, which were situated in open water. In November all samples were from open water, except November station N02, which was in open drift ice (see Wilson et al., 2017, for the definition of the different ice types).

### 2.2 Physical and chemical conditions

#### 2.2.1 Light

At the sampling positions for the January and March cruises, the sun is below the horizon from about mid-October to the beginning of March, and the civil polar night, i.e. when the sun is always more than 6 degrees below the horizon, lasts from the beginning of November until the beginning of February. At the locations sampled during the May, August and November cruises, the sun appears over the horizon from around the 20th February, and there is midnight sun from around the 16th of April. The midnight sun then lasts until the end of August, the sun is again below the horizon from mid-October, and the civil polar night starts around 10th November and lasts until the end of January.

#### 2.2.2 Hydrographical conditions

Vertical profiles of temperature, salinity, and fluorescence were recorded at each sampling station using an SBE 911plus CTD system (Sea-Bird Scientific USA, Bellevue, WA, USA). Hydrographical conditions are described in detail in Ran-delhoff et al. (2018) and data have been deposited at the PANGAEA Data Publisher for Earth and Environmental Science (https://doi.pangaea.de/10.1594/PANGAEA.884255). Briefly, conditions were dominated by the large-scale inflow of warm Atlantic Water (the West Spitsbergen current), which is modified as it enters the cold Arctic Ocean. Surface temperature was highest in August, station P05, ≃ 6 *°*C. Surface temperature and salinity were generally lower at the stations farther off the slope compared to those on the shelf slope (Figure 2). The difference between stations diminished by depth, and at 1000 m the conditions were almost identical across stations and months (Figure 2).

**Figure 2.**
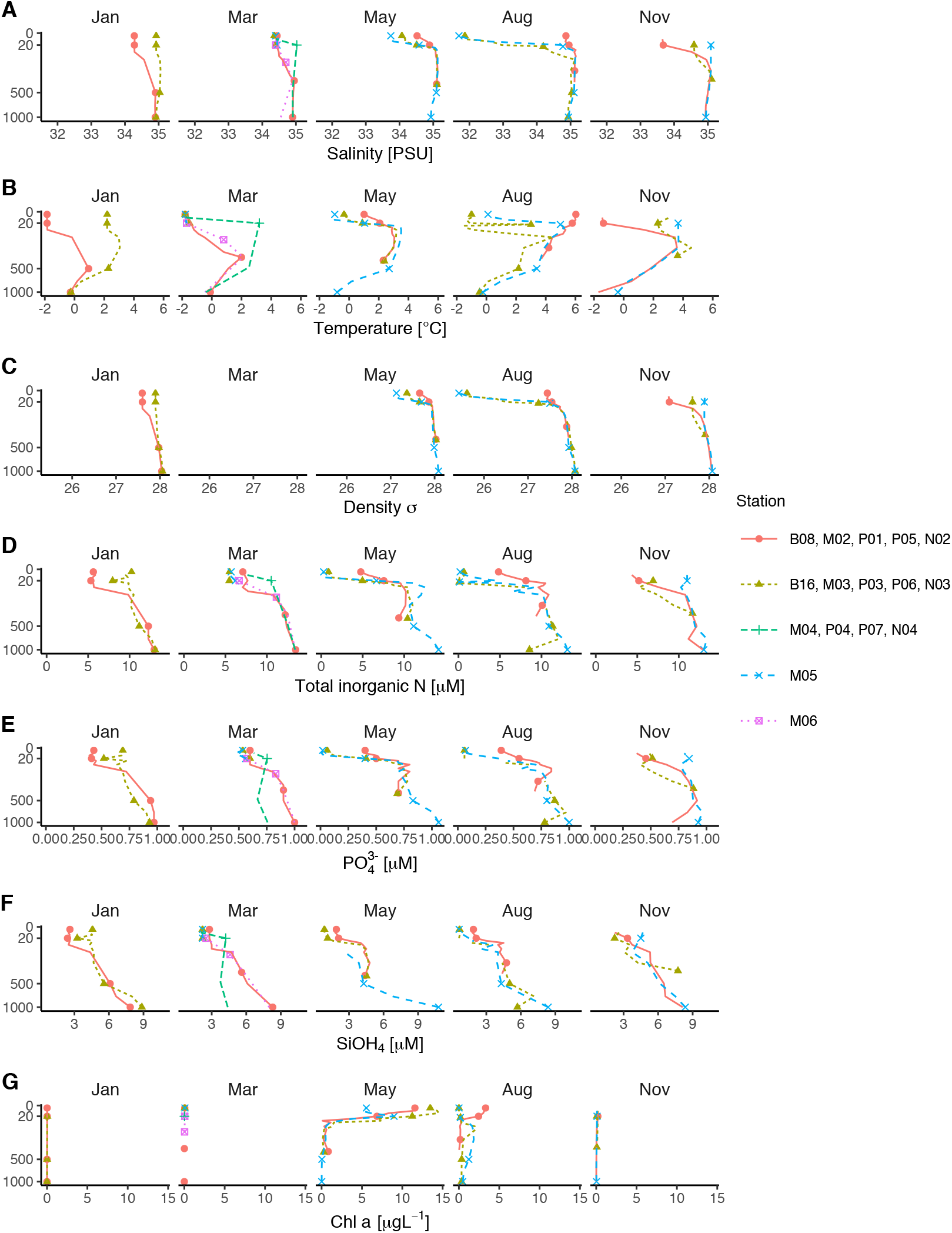
Profiles of environmental variables measured during the MicroPolar cruises. A) Salinity [PSU], B) Temperature [*°*C], C) Density [*σ*], D) Total inorganic N [*µ*M], E) PO_4_^3^ – [*µ*M] F) SiOH_4_ – [*µ*M], G) Chl *a* [*µ*gL^− 1^]. Data obtained from (Paulsen et al., 2017).

#### 2.2.3 Inorganic nutrients and Chlorophyll *a*

Atlantic water was the dominant source of nutrients (as indicated by PO_4_^3–^: NO_3_ ^−^, c.f. Randelhoff et al. (2018)). In the surface, inorganic nutrients and Chl *a* showed opposite patterns. As expected, Chl *a*-concentrations were close to zero in the dark winter months (November and January). In March, there was some daylight, but the water column was not yet stratified, which prevented initiation of the spring bloom. Chl *a* concentrations peaked in May (at most 14 *µ*gL^-1^), concomitantly with depletion of inorganic nutrients. From May to August Chl *a* concentration decreased to < 5 *µ*gL^-1^, while the concentrations of inorganic nutrients were still generally low. By November the concentrations of inorganic nutrients in the epipelagic zone had increased and were again back to the levels observed in January and March.

## 3 Sampling strategy

### 3.1 Sample preparation for DNA extraction

#### 3.1.1 Niskin bottles

From each station and depth, 50 L of seawater were collected in 10L Niskin bottles. The samples were size fractionated. During the January and March cruises, the samples were prefiltered through a 180 *µ*m mesh size nylon filter, and size fractionated into the 3-180 *µ*m and 0.45-3 *µ*m fractions by filtration using a peristaltic pump (Masterflex 07523-80, ColeParmer, IL, USA), through serially connected 3 *µ*m and 0.45 *µ*m polycarbonate filters (Isopore/Durapore, 142 mm diameter, Millipore, Billerica, MA, USA), mounted in stainless steel tripods (Millipore). The filters were removed from the filter holders and cut in four. Two of the pieces were used for DNA extraction, the others were saved for other purposes. The pieces for DNA were transferred to a 50 mL Falcon tube with 1 mL (65 *°*C) AP1 lysis buffer (Qiagen, Hilden, Germany), the plankton material was washed off the filters, and buffer with material and the filters were transferred to two separate cryovials. AP1 buffer (65 *°*C) was added to the vial with the filters, flash frozen in liquid nitrogen and kept at −80 *°*C until DNA extraction. During the May, August and November cruises the water was sequentially filtered through 200, 50, and 10 *µ*m nylon mesh, the material on each nylon mesh was collected with sterile filtered seawater in a 50 mL Falcon tube, and collected by filtration on a polycarbonate filter (10 *µ*m pore size 47 mm diameter, Millipore). The filters were transferred to cryovials to which 1 mL of warm AP1 buffer was added, flash frozen in liquid nitrogen and kept at −80 *°*C until DNA extraction. The size fraction < 10 *µ*m passing through the nylon mesh system was fractionated into the 3-10 *µ*m and 0.45-3 *µ*m size fractions by serial filtration through 142 mm diameter polycarbonate filters as described above.

#### 3.1.2 Net hauls

Vertical phytoplankton net hauls (mesh size 10 *µ*m) were collected between 50 m depth and the surface at each station in May, August and November. The plankton samples were diluted to 1 L with sterile filtered sea water, and size fractionated by filtration through 200, 50 and 10 *µ*m nylon mesh. The plankton was washed off the nylon mesh with sterile sea water, diluted to 50 mL in a Falcon tube and a 20 mL aliquot collected on a 10 *µ*m pore size polycarbonate filter and preserved for DNA extraction as described above. The remaining 30 mL were preserved for microscopical analyses to be reported separately.

### 3.2 DNA extraction

DNA was extracted with the DNeasy Plant mini kit (Qiagen), according to the protocol from the manufacturer, except for the following step: To disrupt the thick cell walls of certain protist groups, the frozen samples in cryovials were incubated at 95 *°*C for 15 min, then shaken in a bead-beater 2x 45-60 s. Subsequently 4 *µ*L RNase was added, and the lysate was incubated on a heating block at 65 *°*C for 15-20 min, with vortexing in-between. Purity and quantity of the extracted DNA was assessed with NanoDrop.

## 4 18S rRNA gene amplicon generation for eukaryotic metabarcoding

### 4.1 PCR amplification

The V4 region of the 18S rRNA gene was amplified with primers 18S TAReuk454FWD1 (5’-CCAGCASCYGCGGTAATTCC-3’) and V4 18S Next.Rev (5’-ACTTTCGTTCTTGATYRATGA-3’) (Piredda et al., 2017). The samples were prepared for Illumina sequencing with a so-called dual-index approach (e.g. Fadrosh et al., 2014), where a 12 bp internal barcode was added to both the forward and reverse amplification primers for the initial amplification. In order to pool several samples into one library preparation, 19 unique barcodes for each direction were used. The internal barcodes were designed to give a balanced distribution of the four bases, following the recommendations of Fadrosh et al. (2014). PCR reactions consisted of 12.5 *µ*L KAPA HiFi HotStart ReadyMix 2x (KAPA Biosystems, Wilmington, MA, USA), 5 *µ*L of each primer (1 *µ*M), 10 ng DNA template and PCR-grade water to a final volume of 25 *µ*L. The PCR was run on an Eppendorf thermocycler (Mastercycler, ep gradient S, Eppendorf), with an initial denaturation step at 95 *°*C for 3 min, followed by 25 cycles of denaturation at 98 °C for 20 s, annealing at 65 *°*C for 60 s and elongation at 72 *°*C for 1.5 min, and a final elongation step at 72 *°*C for 5 min. The reactions were performed in triplicate for each sample and pooled prior to purification and quantification. The length of the PCR products was assessed by gel electrophoresis. In all samples, there was a strong band at about 470 bp, and no other bands (data not shown). The PCR products were purified with AMPure XP beads (Beckman Coulter, Brea, USA) using the standard protocol with elution buffer EB (Qiagen), quantified with a Qubit dsDNA High-Sensitivity kit (Thermo Fisher, Waltham, MA, USA) and pooled in equal concentrations to create nine pools with ca. 19 samples in each. The pools were sent to library preparation at the Norwegian Sequencing Centre (Oslo, Norway) and GATC GmbH (Konstanz, Germany) with the KAPA library amplification kit (Kapa Biosystems). Further quality control of the amplicons were made with Bioanalyzer at the sequencing centres prior to Illumina sequencing. Due to issues with the Illumina MiSeq chemistry in 2015, the sequencing was done with a modified HiSeq protocol on two HiSeq runs at GATC. The HiSeq sequencing runs were spiked with 20% PhiX (viral DNA added as sequencing control). For a few samples, we sequenced separately replicate DNA extractions and replicate PCR runs with 60 *°*C annealing temperature (indicated in Table 1). Samples with low number of reads were re-amplified with 30 cycles (with the original DNA as template) and re-sequenced with Illumina MiSeq at the Norwegian Sequencing Centre. In total we sequenced 199 samples separately (referred to as ‘seq_event’ in Table 1).

**Table 1.**
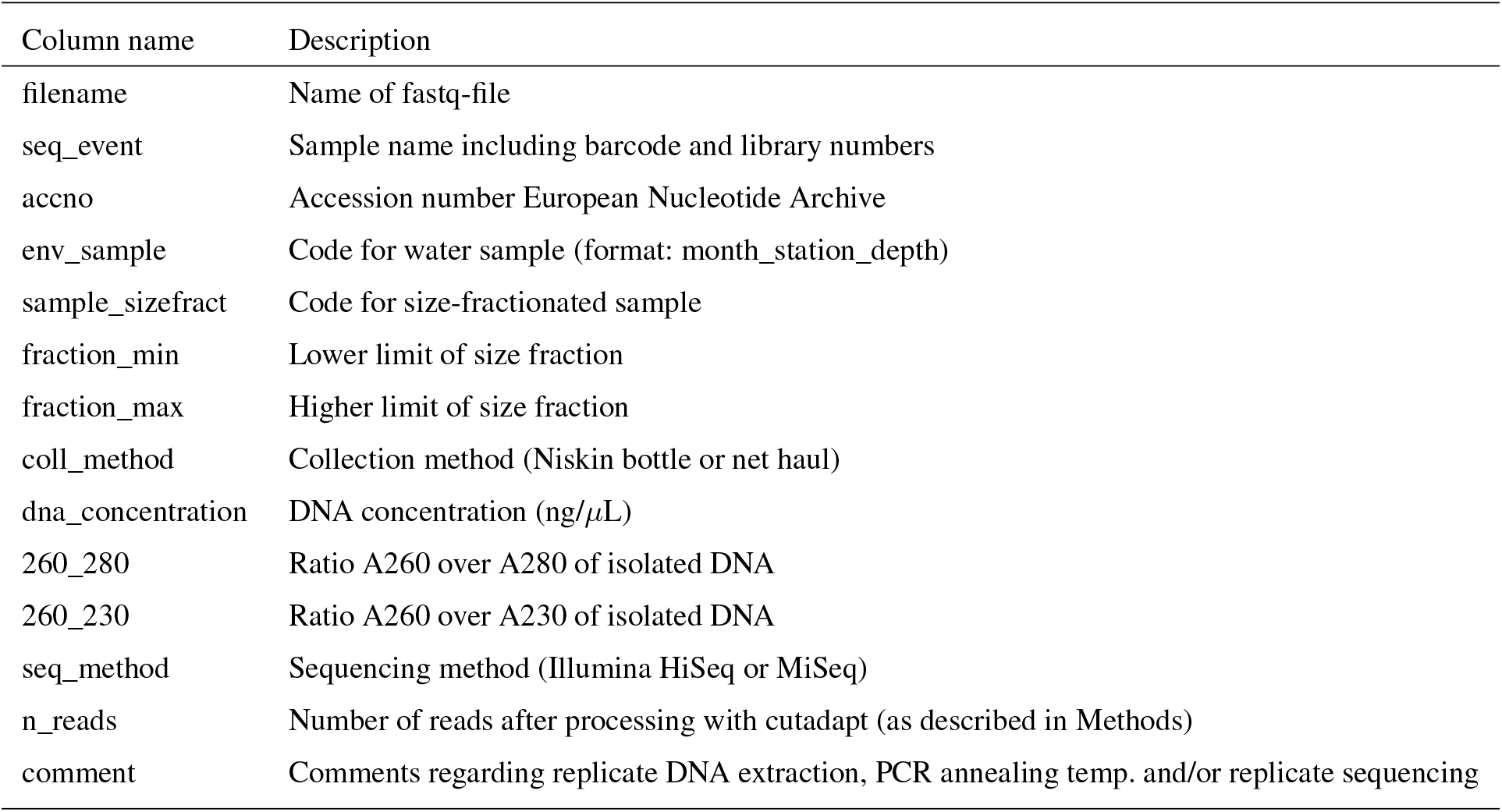
Description of metadata table (named “meta_data_fastqfiles.txt”) for the fastq-files deposited in ENA. These metadata can be joined with environmental data (described in Table 3) by the ‘env_sample’ column. Each fastq-file is unique, but two or more fastq-files may map onto the same DNA-extract and/or PCR.

### 4.2 Bioinformatics processing

PhiX sequences were removed and the raw reads were sorted according to the Illumina index by the Illumina software at the sequencing provider. For the HiSeq datasets, the samples within each Illumina library were demultiplexed with cutadapt v2.10 with Python 3.6.11 (Martin, 2011), requiring 0 errors in the internal barcodes. The amplification primers were removed with cutadapt v2.8 with Python 3.7.6, with setting --trim-n (trim N’s on ends of reads). The reads were denoised and merged with dada2, v1.16. (Callahan et al., 2016). For the HiSeq reads the settings were: truncLen = c(240,200), minLen = c(240,200), truncQ = 2, maxEE = c(10, 10), max_number_asvs = 0. Chimeras were detected with isBimeraDenovo with default settings, and removed with removeBimeraDenovo, with ‘method_chimera’ = “pooled”. For the MiSeq reads truncLen and minLen were set to c(270, 240), the other settings were the same as for HiSeq. The reads were subsequently classified with assignTaxonomy, the dada2 implementation of the naive Bayesian classifier method described in Wang et al. (2007), against the Protist Ribo-somal Reference Database (PR2 Guillou et al., 2013). ASVs with less than 90% bootstrap value at class level and/or which comprised less than 10 reads in total were removed. As this study is focusing on the protists, all reads assigned to Metazoa and Viridiplantae (Embryophyceae) were excluded from the processed ASV tables (Table 2).

**Table 2.** Overview over ASV-tables. Commands for creating ASV tables 1-3 from the original ASV table are found in the script ‘asvtables.R’. From ASV tables 1-3, ASVs assigned to division Metazoa or class Embryophyceae have been removed. ASV tables 1-3 are also available as proportions and presence-absence.

| Name | Description |
| --- | --- |
| metapr2_wide_asv_set_207_208_209_Eukaryota.xlsx | Original ASV table after processing with dada2, including taxonomic classification against PR2. |
| asvtab1_nonmerged_readnum.txt | 'Sequencing events' (i.e. replicates of 'sample_sizefract') kept separate, not subsampled. ASVs assigned to Metazoa and Embryophyceae removed. |
| asvtab2_merged_readnum.txt | Replicates of 'sample_sizefract' merged |
| asvtab3_merged_subsamp_readnum.txt | Replicates of 'sample_sizefract' merged, then all 'sample_sizefract' are subsampled to equal read number within each size fraction. |

### 4.3 Preparation of ASV-tables

Preparation of the ASV-tables was done in R v. 3.6.0 (R Core Team, 2019). Subsampling to equal read number was done with the function rrarefy() from the ‘vegan’ package (Oksanen et al., 2020). Prior to assessing ASV richness, data from fastq-files that map onto the same size-fractionated sample were merged, and all the size-fractionated samples were subsampled as follows: 0.45-3 *µ*m: 40,000 reads, 3-180 *µ*m: 88,000 reads, 3-10 *µ*m: 40,000 reads, 10-50 *µ*m: 40,000 reads, 50-200 *µ*m: 8,000 reads. Subsampling to equal read number was performed 100 times, and the average read number per ASV was used, rounded to 0 decimals. The low number of protist reads in the 50-200 *µ*m fraction was due to a high proportion of Metazoan reads. An R script for merging and sub-sampling is provided in the GitHub repository (scr/asvtables.R), overview of the available versions of the asv-table is given in Table 2, and interactive versions of figures, tables, and supplementary material are available as a Shiny app (Chang et al., 2019) (see section “Data and code availability” below). Figures were made with the R packages ‘ggplot2’ (Wickham, 2016) and ‘plotly’ (Sievert, 2020). Interactive tables were made with the package ‘DT’ (Xie et al., 2020).

## 5 Data description

### 5.1 Overview of sequenced samples

In total we obtained 44 Niskin samples and 8 net hauls, which were fractionated into 140 and 15 size-fractionated samples, respectively (May_P4_net_10_50 failed). These samples are in the following referred to as ‘sample_sizefract’, and de-noted Month_Station_Depth_minfract_maxfract or Month_Station_net_minfract_maxfract. On some ‘sample_sizefract’, we performed replications of DNA extraction, PCR with variable annealing temperature, and/or replicate sequencing. Thus, one or more fastq-file pairs can map onto the same ‘sample_sizefract’. The fastq-files were deposited individually to ENA, and are referred to as a ‘sequencing event’ (’seq_event’). In total the dataset consists of 199 sequencing events, some of which were merged, to form in total 155 ‘sample_sizefract’. Description of metadata available for each fastq-file pair can be found in in Table 1. Table 3 describes all environmental parameters obtained from each water acquisition event (i.e. from the Niskin samples). These are referred to as ‘env_sample’ and labelled Month_Station_Depth.

**Table 3.** Overview of the environmental data available from Paulsen et al. (2017) and Randelhoff et al. (2018). Count data from Sandaa et al. (2018). Can be found in the tables “env_data_depths.txt” and “env_data_profiles.txt”.

| Category | Variables |
| --- | --- |
| Sample name | env_sample (Month_Station_Depth) |
| Time | year-month-day |
| Station name | station |
| Location | latitude (N), longitude (E) |
| Depth (m) | depth_m |
| Physical | temperature, salinity, density |
| Inorganic nutrients | NH<sub>4</sub><sup>+</sup>, NO<sub>2</sub><sup>-</sup>, NO<sub>3</sub><sup>-</sup>, PO<sub>4</sub><sup>-</sup>, SiOH<sub>4</sub><sup>-</sup> |
| Organic compounds (Dissolved, Particulate, Total) | carbon, nitrogen |
| Chlorophyll | total Chl *a*, Chl *a* < 10μm |
| Counts | virus (small, medium, large), het. bacteria, *Synechococcus*,<br>picophytoplankton, nanophytoplankton, *Phaeocystis*, cryptophytes |

### 5.2 Total number of reads and ASVs

After quality filtering, removal of chimeras, singletons and non-target taxonomic groups, the dataset comprised 6536 protist ASVs, corresponding to 32,164,445 reads. After subsampling to equal number of reads per sample within each size fraction, the data set was reduced to 6430 ASVs and 5,729,358 reads. Number of ASVs per division or class within each size fraction, after subsampling, is shown in Table A1. In total we recovered 3,339, 2,720, 2,799, 1,153 and 3,172 ASVs in the 0.45-3, 3-10, 10-50, 50-200 and 3-180 *µ*m size fractions, respectively. Note that the numbers are not directly comparable, as the fractions were not obtained from the same number of samples. Syndiniales and Dinophyceae had the highest number of assigned ASVs, with 2,166 and 1,723, respectively. Ciliophora, Bacillariophyta, Radiolaria and Chlorophyta had between 400 and 200 assigned ASVs each (Table A1).

### 5.3 Sample saturation

Slopes of rarefaction curves at the endpoint, after subsampling, ranged from 0 to 0.014 (Figure 3) which means that for every 1000 extra reads sequenced, we could expect to find between 0 and 14 new ASVs (de Vargas et al., 2015). There was no correlation between the number of ASVs detected in a sample and the slope of the rarefaction curve (r^2^ = −0.13, p = 0.11).

**Figure 3.**
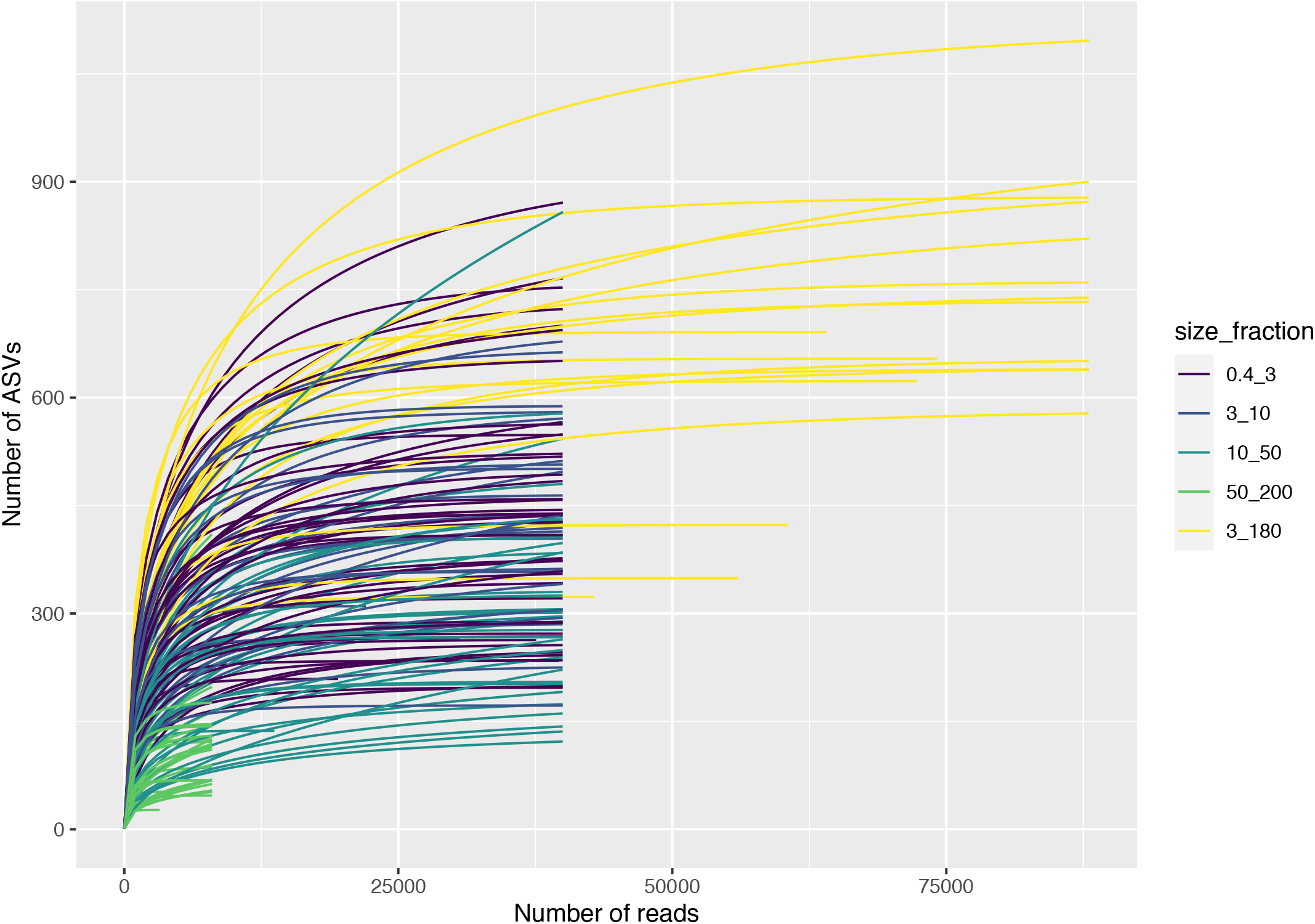
Rarefaction curves for each ‘sample_sizefract’, after subsampling to equal number of reads within each size fraction. Based on “asvtab3_merged_subsamp_readnum.txt”.

### 5.4 Variation in taxonomic composition by season, depth and size fraction

The taxonomic composition of the metabarcoding reads, at division or class levels, is shown in Figure 4. The taxonomic composition of the ASVs in each sample is shown in Figure A1. The metabarcoding data reveal variation in taxonomic composition both by season and depth, in all size fractions. In the following, the fractions are defined as follows: 0.45-3*µ*m = picoplankton, 3-180*µ*m = nano-micro, 3-10*µ*m = nanoplankton, 10-50*µ*m = small microplankton and 50-200*µ*m = large microplankton. All the major protist groups varied from less than 1% of the reads, to up to 99% for the most abundant (e.g. Syndiniales in the picoplankton fraction, and diatoms in the microplankton fractions; Table A2).

**Figure 4.**
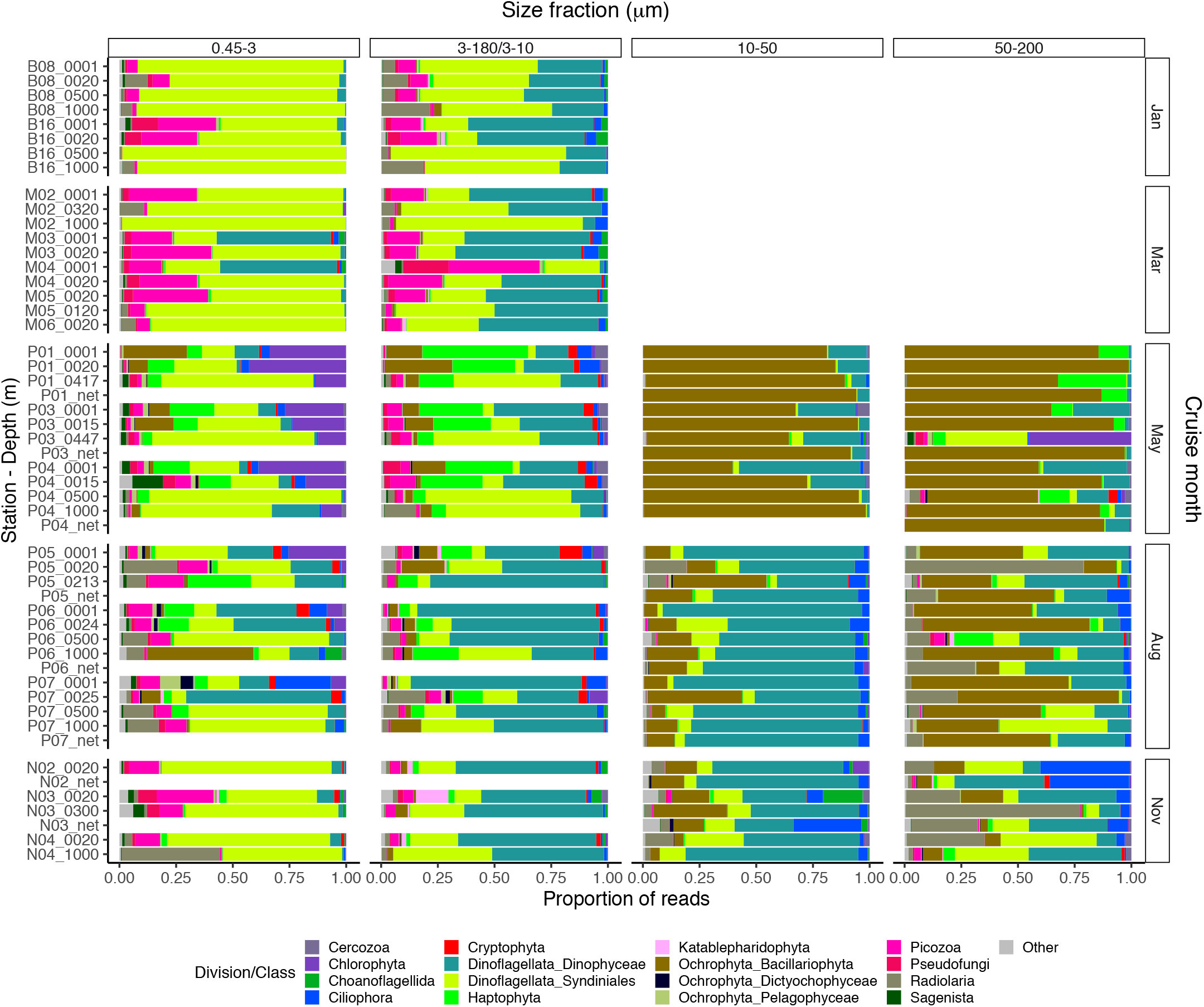
Barplot of relative read abundance of the major protist divisions or classes in each size-fractionated sample. Based on “asvtab3b_merged_subsamp_prop.txt”. Size fraction 3-180/3-10 corresponds to 3-180 *µ*m in January and March, and 3-10 *µ*m otherwise. The 10-50 and 50-200 *µ*m fractions were not available from the January and March cruises. Net hauls were sampled in May, August and November, and were fractionated into the 10-50 and 50-200 *µ*m fractions.

In January (winter) and March (pre-bloom), heterotrophic or parasitic groups were dominating at all depths. In the picoplankton size fraction, the parasitic dinoflagellate group Syndiniales had highest read abundance these months, with up to 99% of the reads, followed by the heterotrophic group Picozoa, with up to 35% of the reads, and Pseudofungi with up to 12% (previously categorised as Marine Stramenopiles, MAST). Syndiniales also had the highest ASV richness in all samples. In the nano-micro fraction Dinophyceae had generally higher read abundance, with 20-55% of the reads in most samples. Syndiniales and Picozoa had up to 82% and 40%, respectively. Syndiniales had highest ASV richness also in this fraction, followed by Dinophyceae. Other heterotrophic groups notably present in this fraction were Pseudofungi and Radiolaria, with 2-20% of the reads each, and Ciliophora and Choanoflagellida with up to 6% of the reads. ASVs assigned to phototrophic groups were detected these months, but constituted less than 3% of the reads in all samples.

The May samples were characterised by higher proportions of phototrophs in all size fractions. In the pico- and nanoplankton fractions, there was a pronounced difference between the epipelagic and mesopelagic samples this month. In the picoplankton fraction, Chlorophyta (mainly represented by the genera *Micromonas* and *Bathycoccus*) had high read abundance in the epipelagic samples with 17-43% of the reads. In the nanoplankton fraction, Haptophyta (mainly represented by the genus *Phaeocystis*) and Dinophyceae were the most abundant groups in the euphotic samples with 25-47% and 14-39% of the reads, respectively. The mesopelagic samples in the pico-nano fractions were characterised by high abundance of Syndiniales, with 47-85% of the reads. In the pico-nano fractions, ASV richness was generally higher in the mesopelagic than in the epipelagic samples. In these fractions, Syndiniales generally had the highest number of ASVs, despite having lower proportional read abundance. In the microplankton fractions diatoms were dominating both in the epi- and mesopelagic samples, with up to 99% of the reads. Dinoflagellates (Dinophyceae) was the second most abundant group in the small microplankton fraction, with up to 50% of the reads. In the large microplankton fraction, *Phaeocystis* was also abundant in certain samples, with up to ca. 30% of the reads. In the net haul samples from May, the diatoms were dominating with up to 97% of the reads. Dinophyceae had 10-11%, and Haptophyta constituted 11% in the large microplankton fraction from station P01. These fractions generally had lower ASV richness than the pico-nano, and there was no clear difference in ASV richness by depth. The groups with highest ASV richness in these samples were Dinophyceae, Bacillariophyta and Syndiniales.

In August, in the picoplankton fraction of the epipelagic samples, Dinophyceae had the highest read abundance, with 20-64%. Haptophyta had ca. 4-14% and Chlorophyta ca. 7-25% in these samples. In the mesopelagic pico-planktonic samples, Syndiniales also dominated in August, with up to 68% of the reads. Radiolaria accounted for 10-13% of the reads in these samples, whereas Picozoa 3-15%. Picozoa relative abundance reached also up to 12% in the epipelagic samples. In the nanoplankton size fraction, Dinophyceae was dominating, with 27-88% of the reads. Similar to in May, Syndiniales had generally highest ASV richness in the picoplanktonic samples. In the nanoplanktonic samples Syndiniales and Dinophyceae had similar ASV richness. In the small microplanktonic fraction, Dinophyceae dominated with 27-88% of the reads. Diatoms constituted up to 41% of the reads, and there was no clear difference in proportion of this group between the epi- and mesopelagic samples. In the large microplanktonic fraction the diatoms dominated, with 30-73% of the reads. Dinophyceae was the second most abundant group in this fraction, with 3-46% of the reads. In the net haul samples, Dinophyceae and diatoms were the most abundant in the microplankton fractions, with 64-77% and 12-20% of the reads, respectively. In the large microplankton fraction Radiolaria were also abundant representing 7-30% of the reads. ASV richness was slightly higher than in May in the microplanktonic fractions. Syndiniales and Dinophyceae had highest richness also in these fractions, followed by diatoms and ciliates.

In November, the proportion of reads assigned to phototrophs was less than 3% in most samples in the pico- and nanoplankton. In the microplankton fractions, diatom reads constituted 1-33%. In the pico fraction, Syndiniales and Picozoa were the most abundant, with 40-75% and up to 25% of the reads, respectively. Radiolaria represented 44% of the reads in the sample N04_1000. Dinophyceae was the most abundant group in the fractions between 3 and 50*µ*m, with 28-76% of the reads. In the large microplankton fraction, Radiolaria, Dinophyceae and Syndiniales were the most abundant with up to 77, 43 and 42% of the reads, respectively. In the net hauls, Ciliophora was also abundant, with up to ca. 30% of the reads in each size fraction. ASV richness was generally higher this month than in May and August, especially in the nano and small microplanktonic fractions. Syndiniales and Dinophyceae had highest richness also this month.

## 6 Conclusions

This dataset offers novel insights into the spatial and seasonal diversity and dynamics of the protist community in the Atlantic gateway to the Arctic Ocean. It is the first study to provide data on the eukaryote microbial food web throughout a complete year and down to 1000 m in this area of the Arctic. It forms the basis for future studies to detect changes in the eukaryote microbial community, and for more detailed studies on the dynamics and community structure of specific taxonomic groups.

## Code and data availability

The fastq files with raw 18s rDNA V4 reads are available on the European Nucleotide Archive repository under project number PRJEB40133. The original ASV table, meta data table and a table with environmental data obtained from water samples corresponding to the size-fractionated plankton samples are deposited in the Sea scientific open data publication repository (SEANOE), with doi: https://doi.org/10.17882/79823. The ASV tables, including the ASV sequences and assigned taxonomy, R-scripts for producing the figures and tables, and a Shiny application with interactive versions of the figures and tables are deposited on GitHub: https://github.com/EEgge/micropolar_protists_datapaper. The Shiny app can be opened in RStudio by running the following command: shiny::runGitHub(“micropolar_protists_datapaper”,”EEgge”, ref = “main”).

## Author contributions

Conceptualization: AL, GB, BE, DV, UJ, Data curation: EE, BE, DV, Formal analysis: EE, BE, DV, Funding acquisition: AL, GB, BE, DV, UJ, Investigation: AL, GB, BE, UJ, SE, EE, Project administration: AL, GB; Visualization: EE, DV, BE; Writing - original draft preparation: EE, BE, DV; Writing - review and editing: EE, BE, DV, AL, GB, SE, UJ. All authors read and approved the final version of the paper.

## Acknowledgements

The study was conducted as part of the Research Council of Norway supported project Micropolar - Processes and Players in Arctic Marine Pelagic Food Webs - Biogeochemistry, Environment and Climate Change no. 225956/E10 (prosjektbanken.forskningsradet.no/#/project/NFR/225956/). DV was supported by ANR contract PhytoPol (ANR-15-CE02-0007). We wish to thank members of the MicroPolar and Carbon Bridge projects for assisting in the sampling campaigns, and the crews at K/V Svalbard (January cruise), R/V Lance (March) and R/V Helmer Hanssen (May, August and November cruise).

## Appendix A

### Supplementary material

**Figure A1.**
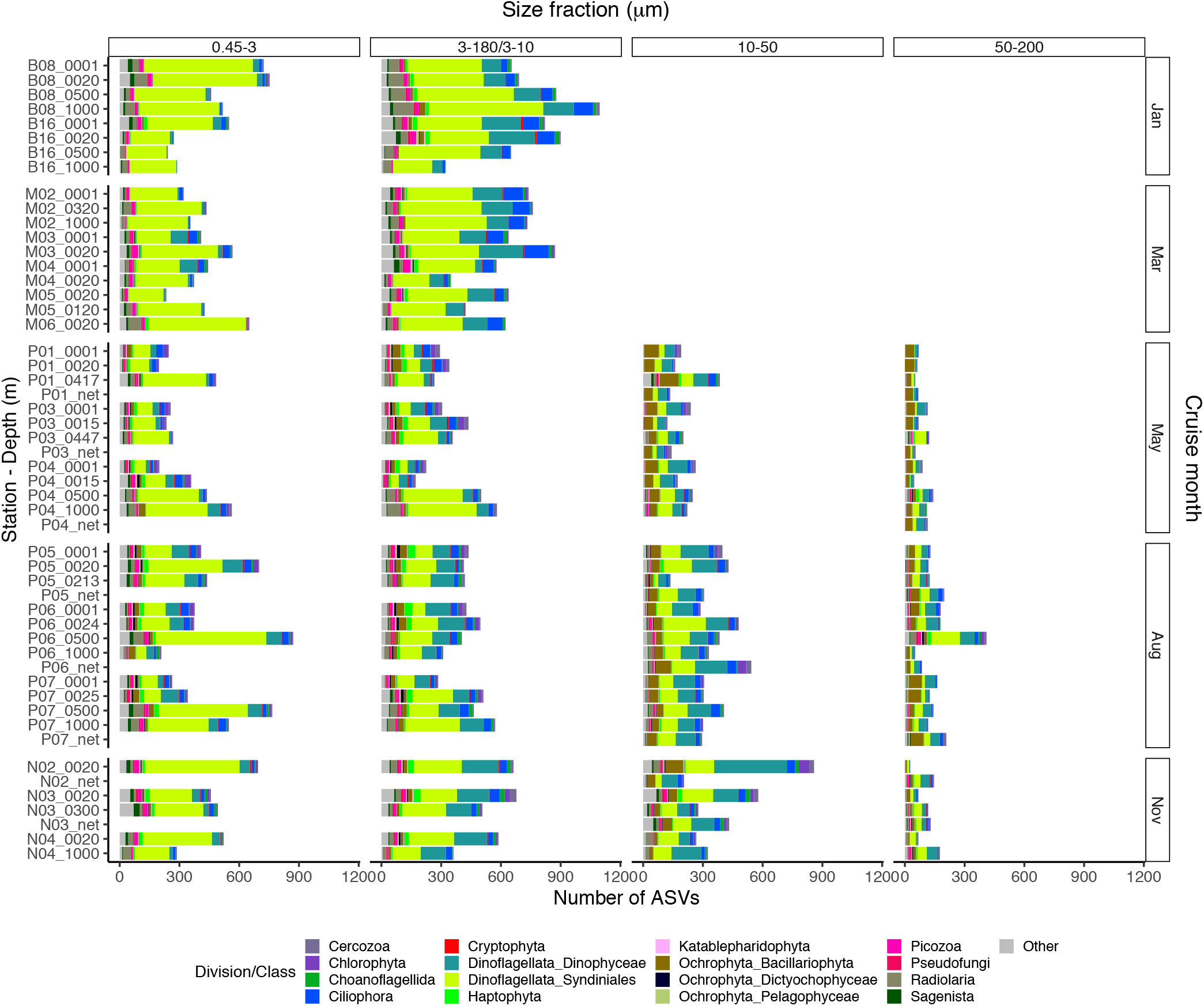
Barplot of number of ASVs of the major protist divisions or classes within each ‘sample_sizefract’. Based on “asvtab3c_merged_subsamp_pa.txt”. Size fraction 3-180/3-10 corresponds to 3-180 *µ*m in January and March, and 3-10 *µ*m otherwise. The 10-50 and 50-200 *µ*m fractions were not available from the January and March cruises. Net hauls were sampled in May, August and November, and were fractionated into the 10-50 and 50-200 *µ*m fractions.

**Table A1:**
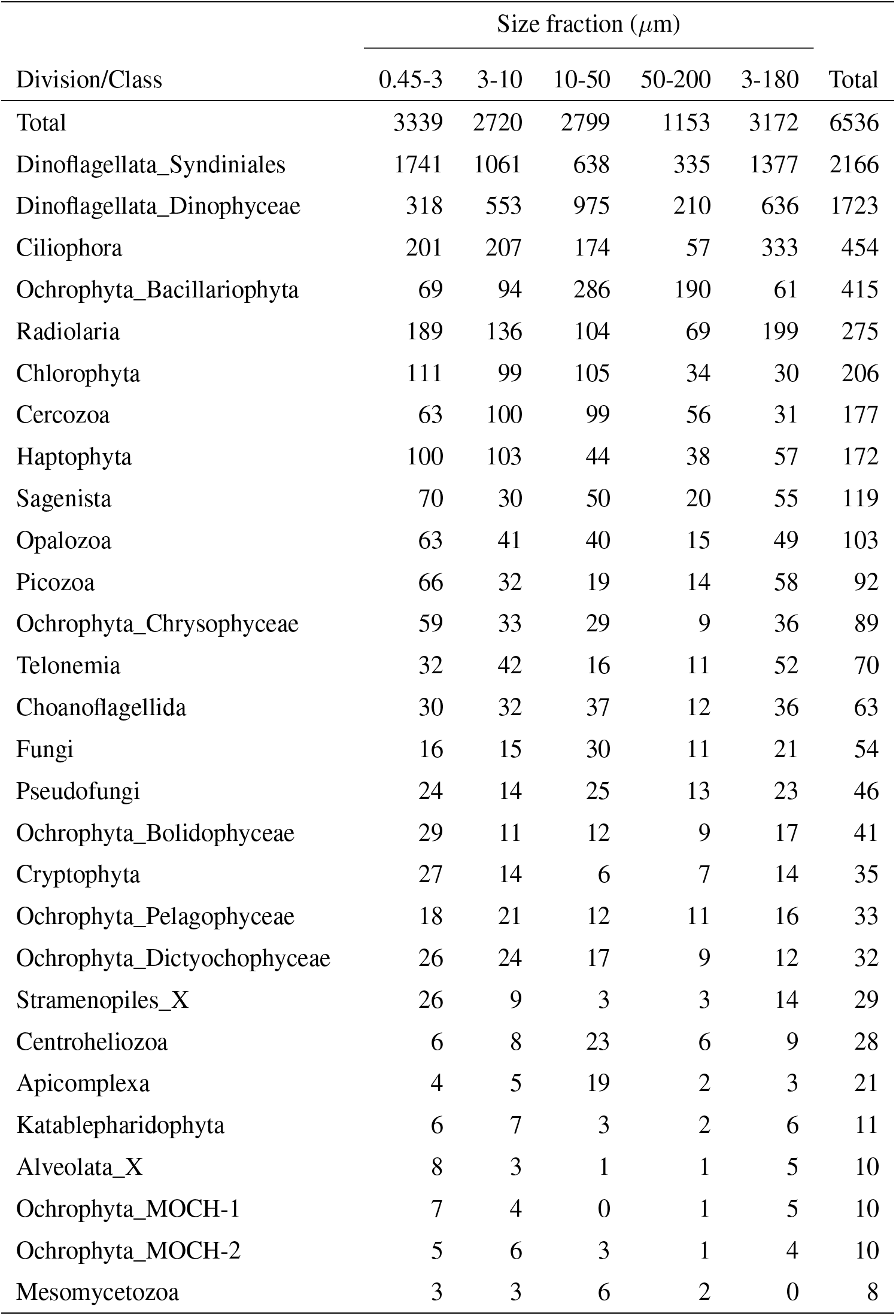
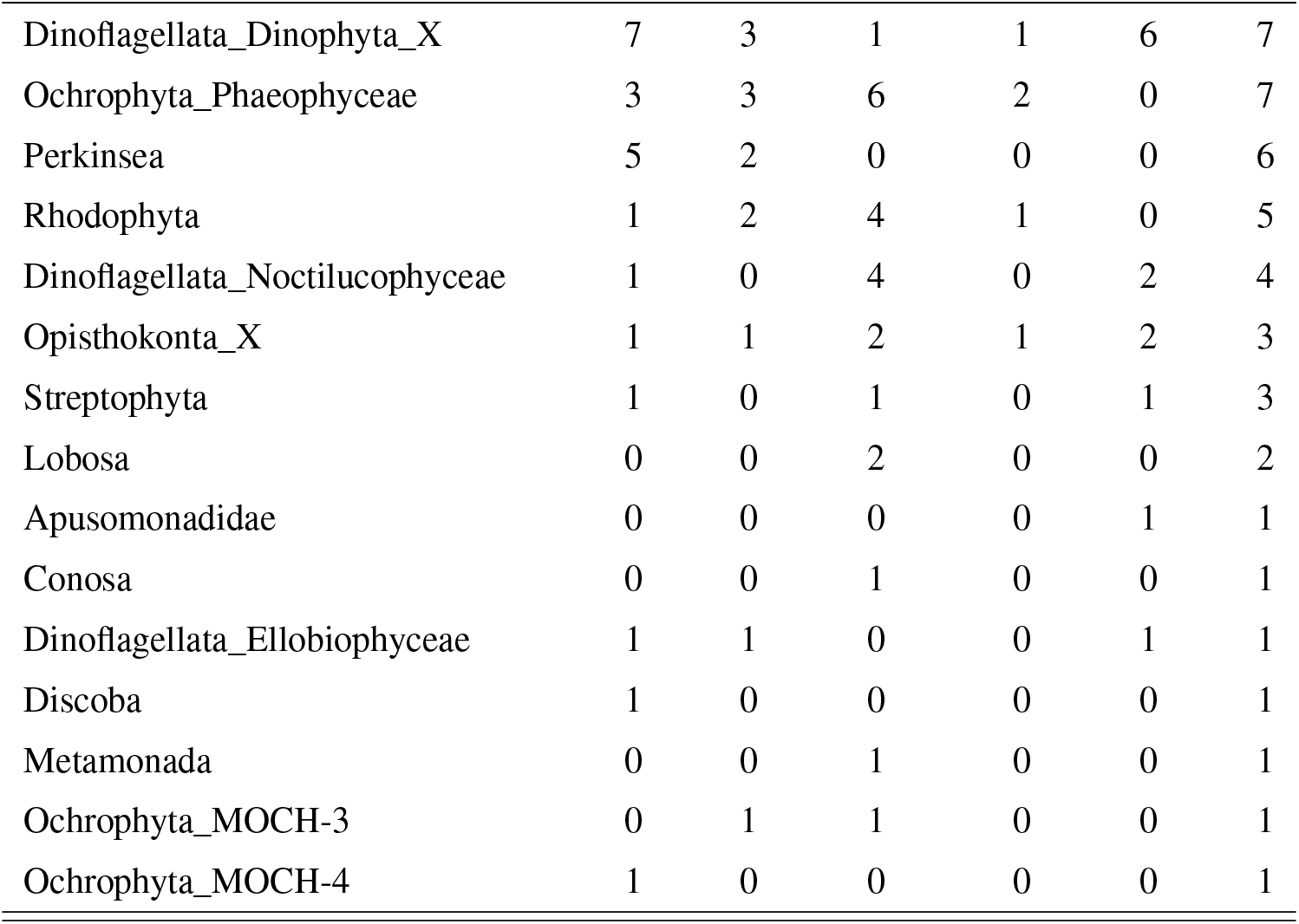
Number of ASVs assigned to each division or class, distributed by size fraction, and in total. Note that a given ASV may occur in multiple size fractions.

**Table A2:**
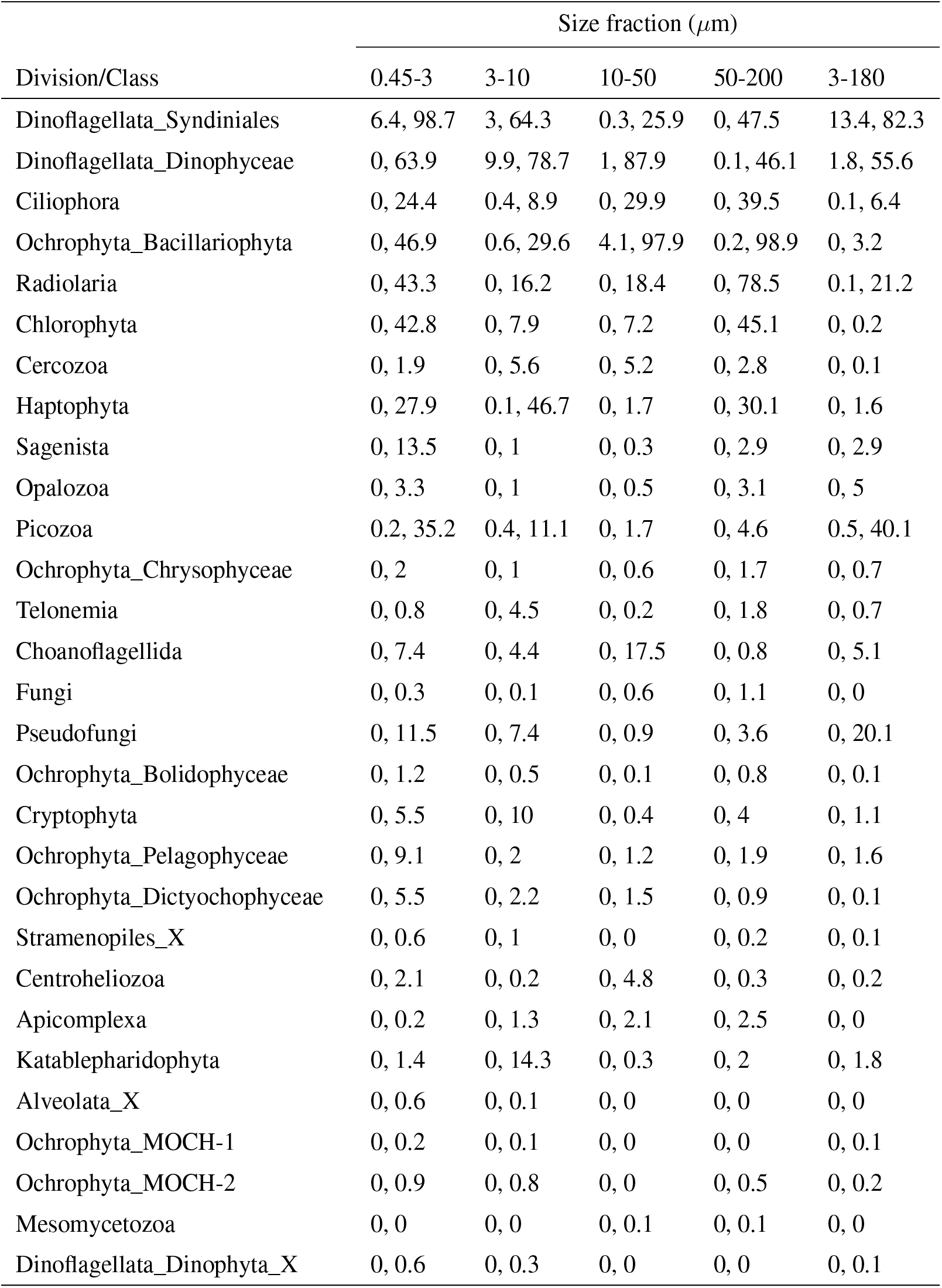
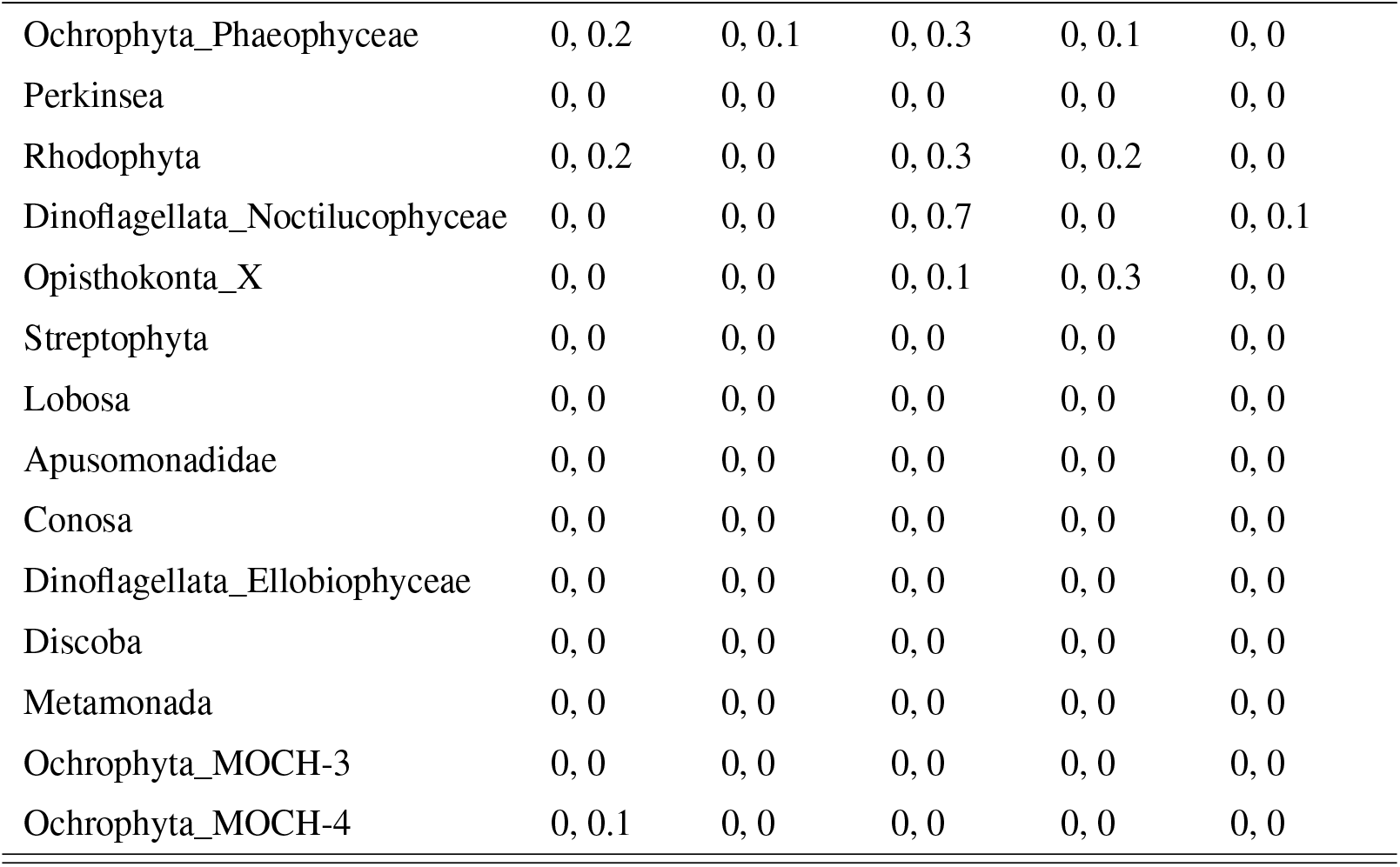
Minimum and maximum percentage of reads of each division or class in each size fraction. The entries have the format ‘min, max’

